# CtBP restricts DUX-dependent and -independent genetic program for the 2-cell-like state in murine embryonic stem cells

**DOI:** 10.1101/2024.01.12.575352

**Authors:** Kazuma Yoshioka, Kota Sugiyama, Maki Ichisakino, Selma Alamanda Abadi, Nao Hayakawa, Miyu Marutani, Ryo Masuda, Kyo Takahashi, Yoshiyuki Seki

## Abstract

After fertilization, maternally deposited mRNA is cleared, and *de novo* mRNA is transcribed from the zygotic genome through zygotic genome activation (ZGA), a process known as maternal-to-zygotic transition (MZT) occurring in the mouse at 2-cell (2C) stage. 2C-like cells (2CLCs) marked by MERVL expression are transcriptionally similar to 2C embryos spontaneously emerge from mouse embryonic stem cells (mESCs). Although the emergence of 2CLCs completely depends on DUX function, a recent knockout study clearly showed that DUX is dispensable for mouse embryos, suggesting that DUX-independent molecular pathways are not recapitulated in 2CLCs. We present here that the disruption of C-terminal binding protein 1/2 (*Ctbp1/2*) activates DUX-dependent and -independent molecular pathway associated with the development of early mouse embryos mediated by the upregulation of Preferentially expressed antigen of melanoma family-like 7 (PRAMEL7). Furthermore, the abnormality of the gene expression profile caused by *Dux* KO is partially rescued by the overexpression of PRAMEL7 in mESCs. Our study provides new insights into the DUX-independent molecular pathway for the activation of early embryonic genes in mESCs.

## Introduction

After fertilization, maternal factors direct development and trigger zygotic genome activation (ZGA) at the around late one-cell stage in mice, called maternal-to-zygotic transition (MZT)(Eckersley-Maslin et al., 2018). The early-cleavage stage embryos acquire cellular totipotency to generate both embryonic proper and extraembryonic tissues during MZT. Although totipotency was thought to be exclusive to the early-cleavage stage embryos, a part of the molecular network is recapitulated in the totipotent-like cells (2-cell-like cells; 2CLCs) marked by the expression of endogenous retrovirus MERVL, which are an endogenous subpopulation of mouse embryonic stem cells (ESCs). (Macfarlan et al., 2012a) (Rodriguez-Terrones et al., 2018).

The transition from pluripotent stem cells into 2CLCs is a useful *in vitro* system for identifying the gene regulatory network for ZGA, and several genes that control the emergence of 2CLCs from pluripotent cells have been identified (Eckersley-Maslin et al., 2019; Hu et al., 2020; Ishiuchi et al., 2015; Percharde et al., 2018; Rodriguez-Terrones et al., 2018). The transcription factor DUX (DUX4 in humans) is a critical factor for regulating the transition from pluripotent stem cells into 2CLCs and the expression of ZGA genes (De Iaco et al., 2017; Hendrickson et al., 2017; Whiddon et al., 2017). Ectopic expression of DUX enhances the population of 2CLCs, whereas the *Dux* disruption eliminates the emergence of spontaneous 2CLCs during mESC culture. Several 2CLC regulatory genes have been identified, but they are either direct activators (DPPA2/4, p53 and NELFA) or direct repressors (CAF-1, LINE-1 and SETDB1) of *Dux* expression (Eckersley-Maslin et al., 2019; Grow et al., 2021; Ishiuchi et al., 2015; Percharde et al., 2018; Wu et al., 2020). However, both *Dux* zygotic and maternal knockout embryos can survive full-term (Chen and Zhang, 2019; Guo et al., 2019), suggesting that DUX-independent molecular pathways not recapitulated in 2CLCs are critical for ZGA and the early development of mice.

In this study, we show that disruption of Ctbp1/2 (C-terminal binding protein 1/2) (Stankiewicz et al., 2014)activates two distinct sets of genes in ESCs: the first set is transiently upregulated in a DUX-dependent manner during minor ZGA, and the second set contains maternally expressed genes and the upregulated genes in a DUX-independent manner during major ZGA. Preferentially Expressed Antigen in Melanoma-like 7 (PRAMEL7) is directly upregulated by the loss of *Ctbp1/2* without DUX function, and upregulated PRAMEL7 activates minor ZGA genes in a DUX-dependent manner and maternally expressed genes and major ZGA in DUX-independent manner in mESCs. These data revealed that CtBP restricts the DUX-dependent and -independent genetic programs for the 2-cell-like state in mESCs.

## Material and Methods

### Cell Culture

E14tg2a ESCs (RIKEN Cell Bank) were cultured in Glasgow minimum essential medium (GMEM; Wako) containing 10% fetal calf serum (FCS; Invitrogen), 1 mM glutamine (Wako), nonessential amino acids (Wako) and 500 μM Monothioglycerol Solution (Wako). The culture was supplemented with 1000 U/ml of leukaemia inhibitory factor (LIF) (Wako) without feeder cells. ESCs were cultured in N2B27 medium with 2i (PD0325901, 0.4 μM: WAKO, CHIR99021, 3 μM: Sigma-Aldrich).

### Generation of knockout ESCs using a CRISPR-Cas9 system

Guide RNAs for *Ctbp1/2*, *Dux* and *Pramel7* were generated in pX330-U6-Chimeric_BB-CBh-hSpCas9 (Addgene plasmid #42230; deposited by Feng Zhang; Cong et al., 2013). The guide RNA sequences are listed in Supplementary Table 1. The pX330 was co-transfected with pCAG-EGFP into E14tg2a with MERVL::TurboGFP (Addgene plasmid #69072; deposited by Maria-Elena Torres-Padilla; (Ishiuchi et al., 2015). The GFP-positive fraction was sorted by Cell Sorter (No. SH800; Sony Biotechnology, Inc., Tokyo, Japan). Genomic DNA deletions were sequenced (Supplementary Figure 1).

### qRT-PCR and RNA-Seq

Total mRNA was purified with a PureLink^TM^ RNA Mini Kit (Ambion, Inc.). For quantitative RT-PCR, cDNA was synthesized by ReverTra Ace® qPCR RT master Mix (Toyobo), and then analyzed by qPCR with FastStart SYBR Green Master Mix (Roche) using a specific primer set for each gene (Supplementary Table 1). RNA-seq libraries were generated using Collibri^TM^ Stranded RNA library prep for ilIumina^TM^ Systems (ThermoFisher Scientific) and sequenced using HiSeq X^TM^ Ten (Illumina) (150-bp paired-ends reads).

### Data analysis of RNA-Seq and public ChIP-seq data

Total mRNA of each cell line was purified with a PureLink^TM^ RNA Mini Kit (Ambion, Inc.) by biological duplicate. RNA-seq libraries were generated using Collibri^TM^ Stranded RNA library prep for ilIumina^TM^ Systems (ThermoFisher Scientific) and sequenced using HiSeq X^TM^ Ten (Illumina). Reads were aligned by Hisat2 to the Mus musculus genome (mm10). Raw read counts of genes were calculated by R package FeatureCounts using mm10 KnownGenes. GTF from UCSC for annotation. Normalized count data, principal component and K-means clustering analysis were exported by the interactive Expression and Pathway analysis (iDEP) webserver using the count data or normalized expression data of early mouse embryos (Ge et al., 2018). Differentially expressed genes (DEGs) were selected using a cut-off at a false discovery rate (FDR) ≦ 0.1 and a ≧ 2-fold change in expression. Gene set enrichment analysis (GSEA) was used to determine whether upregulated genes by *Ctbp1/2* knockouts were enriched for the upregulated genes in 2-cell embryos or 2CLCs as described previously. Venn diagrams showing overlapping genes were constructed by BioVenn, a web application (Hulsen et al., 2008).

Pubic ChIP-seq data for CtBP2 were re-analyzing using GREAT (Genomic Regions Enrichment of Annotations tool) with the “two nearest genes” setting, which assigns each peak to the two closest transcription start sites (TSSs) within 1 Mb. Peak visualization of ChIP-seq peaks for CtBP2, H3K27Ac and DUX was performed using Integrative Genomics Viewer (IGV) (Hendrickson et al., 2017; Kim et al., 2015; McLean et al., 2010; Zhang et al., 2020).

### Immunofluorescence analysis

Cells were fixed by 4% paraformaldehyde at 4℃ for 20 min, followed by incubation with PBS containing 0.5% Triton-X at 4℃ for 5 min. Permeabilized cells were incubated with a primary antibody solution containing anti-GFP (Nacarai Tesque, 04404-84, 1:500) with anti-ZSCAN4 (Merk Millipore, AB4340, 1:1000) or anti-MuERVL-Gag (Epigentek, A-2801-050, 1:250) or anti-OCT4 (Santa Cruz, Sc-8628, 1:500) or anti-CtBP2 (BE Biosciences, 612044, 1:500) followed by secondary antibody solution containing anti-Rat IgG Dylight^TM^ 488 antibody (Rockland, 612-141-120, 1:500) or anti-Rabbit IgG Dylight^TM^ 549 antibody (KPL, 042-04-15-06, 1:500) or anti-Mouse IgG Dylight® 550 antibody (Abcam, ab96876, 1:500) or anti-Goat IgG Dylight^TM^ 549 antibody (Rockland, 605-742-002, 1:500) with 1 μg/ml of DAPI solution (DOJINDO, 340-07971).

### Luciferase assay

The enhancer candidate region of *Pramel7* or *Dux* was cloned into pGL4.23, driving the *luciferase* gene. Wild-type or *Ctbp1/2* DKO ESCs were transfected with the reported constructs with *Ctbp2* or *Pramel7* expression vector using HilyMax (DOJINDO). Cells were harvested 36 hours after transfection, and the luciferase activities were measured using the Dual-luciferase Reporter Assay System (Promega).

### Western blotting analysis

Cells were lysed by boiling SDS sample buffer (WAKO). Before lysed proteins were applied to polyacrylamide-SDS gels, 2-mercaptoethanol was added to denature the proteins. The lysed proteins were separated on polyacrylamide-SDS gels, blotted on a polyvinylidene fluoride membrane, and probed using the following primary antibodies: anti-UHRF1 (MBL, D289-3, 1:1000) or anti-TUBLIN (SIGMA, T5168, 1:5000). Following the primary antibody reaction, the membrane were incubated with horseradish peroxidase (HRP)-coupled secondary antibodies. All antibodies were detected with Immunostar LD (WAKO).

### Phylogenetic tree inference

Amino acid sequences of PRAME family proteins were obtained from Ensemble release 112. Multiple-sequence alignment was performed using the ClustalW algorithm with the MEGA software, version 11 with default parameters. Molecular phylogenetic trees were inferred with the maximum-likelihood method using MEGA software, version 11 with default parameters.

## Results

### CtBP1/2 acts as negative regulators of the transition from pluripotent ESCs to two-cell-like cells (2CLCs)

We have previously shown that CtBP1/2 are essential for the transition from formative to naïve pluripotency in mouse embryonic stem cells (mESCs) (Yamamoto et al., 2020). To investigate their roles of CtBP1/2 beyond pluripotency maintenance, we performed RNA sequencing (RNA-seq) to compare the gene expression profiles between wild-type (WT) and *Ctbp1/2* double-knockout (DKO) ESCs. An MA-plot revealed a marked upregulation of two-cell (2C)-specific genes, including *Dux*, *Zscan4* and *Zfp352*, together with repeat elements such as MERVL, MT2 and GSAT_Mm in *Ctbp1/2* DKO ESCs (Figure 1A, B). Gene set enrichment analysis (GSEA) using a reference set of 2C embryo and 2C-like cell (2CLC)-enriched genes (n=404; (Macfarlan et al., 2012)) confirmed a strong enrichment of 2C-related transcripts among those upregulated in the absence of CtBP1/2 (Figure 1C).

**Figure 1.**
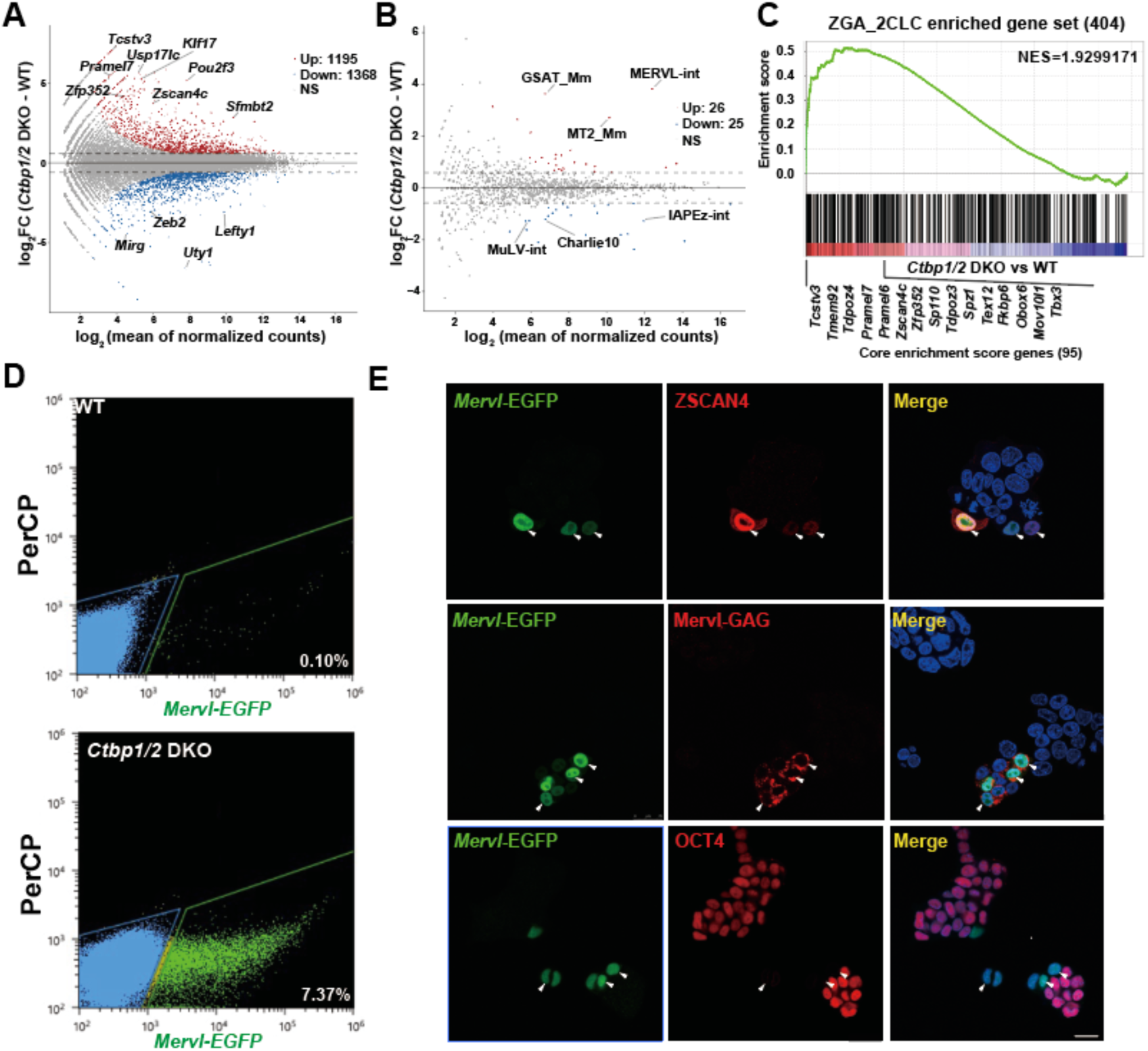
Disruption of *Ctbp1/2* expands the 2-cell-like cell (2CLC) population. (A), (B) MA-plots showing the relationship between average gene (or repeat) expression and log_2_ fold change between *Ctbp1/2* DKO and wild-type (WT) ESCs. Differentially expression was defined as adjusted *p* ≤ 0.05 and fold change ≥ 2. Red and blue dots indicate up- and down-regulated genes, respectively. (C) Gene set enrichment analysis (GSEA) of genes upregulated in 2-cell embryos compared with oocytes and enriched in MERVL-positives 2CLCs relative to MERVL-negative ESCs (GSE33923) (Macfarlan et al., 2012). (D) Flow cytometry analysis of MERVL::TurboGFP reporter activity in WT ESCs and *Ctbp1/2* DKO ESCs. (E) Immunofluorescence analysis showing that MERVL::TurboGFP-positive cells co-express ZSCAN4 and MERVL-GAG proteins, and are mutually exclusive with OCT4 expression. Nuclei were counterstained with DAPI. Scale bar: 10 μm.

To directly monitor the induction of 2CLC state, we generated *Ctbp1/2* DKO ESCs carrying the MERVL-EGFP reporter (Ishiuchi et al., 2015). The GFP-positive fraction was approximately 0.1 % in WT ESC, whereas *Ctbp1/2* DKO ESCs exhibited an over 50-fold increase (7.4%) in GFP-positive cells, indicative of robust 2CLC induction (Figure 1D). These GFP-positive cells co-expressed hallmark 2CLC markers such as ZSCAN4 and MERVL-GAG, while pluripotency-associated factors such as OCT4 were markedly reduced (Figure 1E), consistent with previous reports (Eckersley-Maslin et al., 2016; Zhu et al., 2021).

### CtBP1/2 repress DUX-dependent minor ZGA genes and DUX-independent maternal and major ZGA genes in ESCs

DUX, the homeobox transcription factor, is both essential and sufficient for the emergence of spontaneous 2CLCs from ESCs (De Iaco et al., 2017; Whiddon et al., 2017). To determine whether the 2C-like program triggered by *Ctbp1/2* deficiency depends on DUX, we deleted the *Dux* cluster in both WT and *Ctbp1/2* DKO ESCs (Supplementary Figure 1C). We first compared the population of MERVL-EGFP-positive cells among WT, *Dux* KO, *Ctbp1/2* DKO and *Ctbp1/2*/*Dux* triple-KO (TKO) ESCs. While the strong GFP-positive population was markedly reduced in TKO compared to DKO ESCs, a weak GFP-positive population persisted in the TKO (Figure 2A, B). RT-qPCR analysis confirmed that *Pramel7*, which is upregulated at the 2C stage in WT and *Dux* KO embryos (Guo et al., 2019), remained elevated in both DKO and TKO ESCs, indicating DUX-independent activation (Figure 2C). In contrast, *Zscan4c* upregulation upon *Ctbp1/2* depletion was completely abolished by *Dux* deletion, indicating strict DUX dependency.

**Figure 2.**
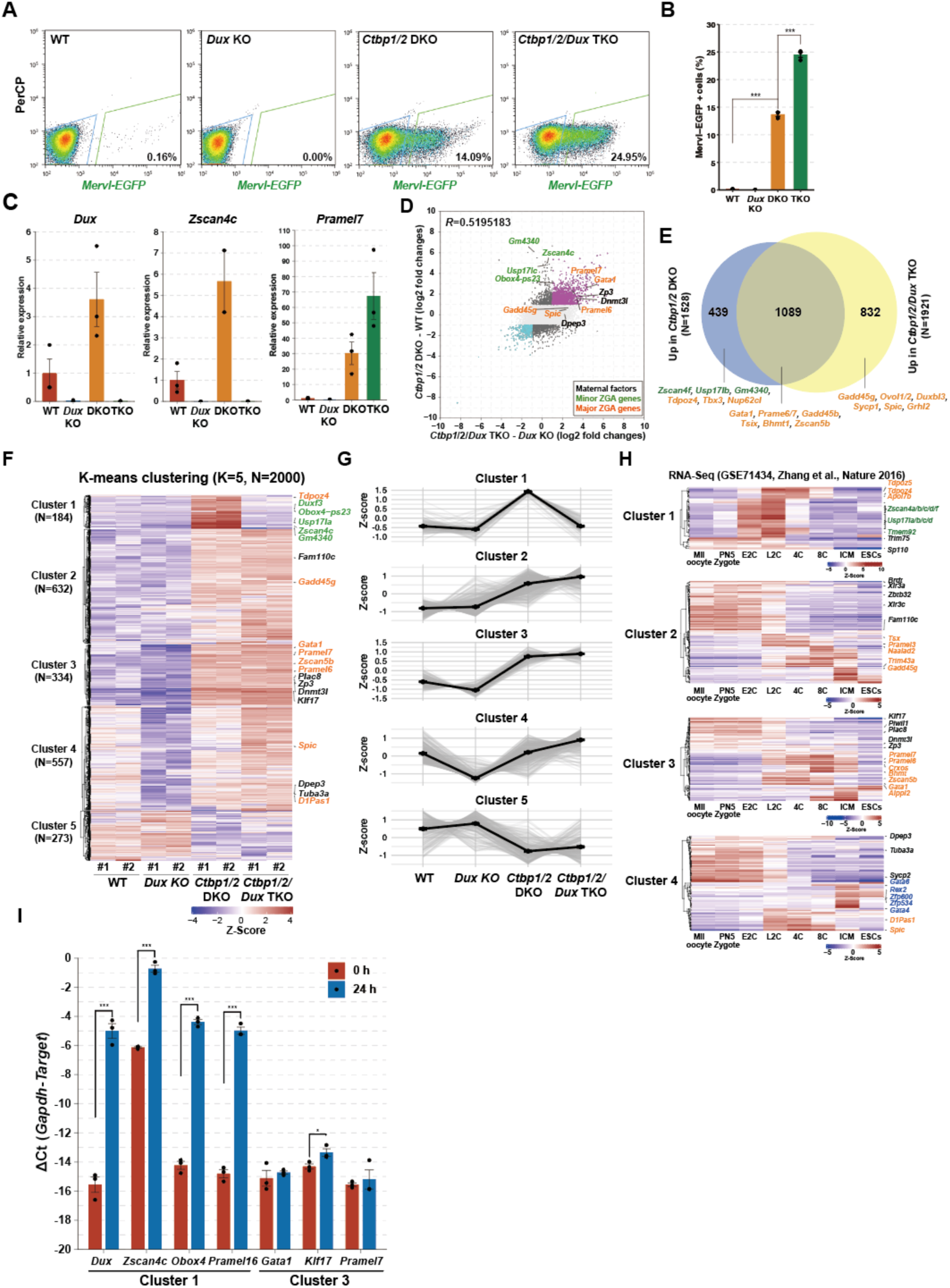
CtBP1/2 repress both DUX-dependent and -independent transcriptional programs associated with preimplantation embryos in mouse ESCs. (A) Flow cytometry analysis (FACS) of MERVL::TurboGFP reporter expression in wild-type (WT), *Dux* knockout (KO), *Ctbp1/2* double knockout (DKO) and *Ctbp1/2*/*Dux* triple knockout (TKO) ESCs. (B) Percentage of MERVL-EGFP-positive cells obtained by FACS Shown are standard error of means (SEM) at biological triplicates. (C) RT-qPCR analysis of 2C-associated genes in WT, *Dux* KO, *Ctbp1/2* DKO and *Ctbp1/2*/*Dux* TKO ESCs. Values are shown as relative expression with WT set to 1. Error bar indicates the standard error of means. \*\*\**P* < 0.001, \*\**P* < 0.005, \**P* < 0.05 (Tukey-Kramer multiple comparison test). (D) Scatter plot comparing gene expression changes induced by *Ctbp1/2* DKO in WT (Y-axis) and *Dux* KO (X-axis) backgrounds. (E) Venn diagram showing the overlap of genes upregulated in *Ctbp1/2* DKO and TKO. (F) K-means clustering analysis (K=5) of the top 2,000 differentially expressed genes across WT, *Dux* KO, *Ctbp1/2* DKO and *Ctbp1/2/Dux* TKO ESCs. (G) Average expression profiles of genes from each cluster at WT, *Dux* KO, *Ctbp1/2* DKO and *Ctbp1/2/Dux* TKO ESCs. Grey lines indicate individual gene, and black line represents the average expression of genes within each cluster. (H) Hierarchical clustering of genes from each cluster using public RNA-seq data of early mouse embryo (Zhang et al., 2016). (I) RT-qPCR after *Dux* induction (0 h vs 24 h) for selected genes (Cluster 1: *Dux, Zscan4c, Obox4, Pramel16 and Cluster 3: Gata1, Klf17, Pramel7*). The ΔCt of each gene was calculated by subtracting the average Ct of Gapdh from that of each gene Ct. Means of ΔCt were calculated from three biological triplicates. Error bar indicates the standard error of means (SEM). \*\*\**P* < 0.001, \*\**P* < 0.005, \**P* < 0.05 (Tukey-Kramer multiple comparison test).

To assess transcriptional activity of the MERVL long terminal repeat (MERVL-LTR) sequences, we performed a luciferase reporter assay (Supplemental Figure 2A). *Ctbp1/2* loss led to robust activation of MERVL-LTR reporter, and mutation of DUX-binding motif (DUXmut-MERVL-LTR) markedlyreduced, but did not abolish, this activity. Reporter activity in *Ctbp1/2/Dux* TKO remained substantially higher than *Dux* KO, indicating that CtBP1/2 also repress MERVL through DUX-independent mechanisms. Published DUX ChIP-seq data (Hendrickson et al., 2017) revealed DUX occupancy near both the *Zscan4c* and *Naalad2* loci, with strong binding signals overlapping MERVL-LTR or MT2 family elements (Supplementary Figure 2B). At the *Zscan4c* locus, DUX binds both proximal regulatory region with a strong peak and distal MERVL elements. At the *Naalad2* locus, DUX-binding sites are exclusively overlapped with MERVL sequences. Reporter assay confirmed that MERVL-independent elements proximal to the *Zscan4c* locus were activated by *Ctbp1/2* loss in a DUX-dependent manner, whereas MERVL-LTR-containing elements derived from the *Zscan4c* and *Naalad2* loci retained partial activity even in *Ctbp1/2/Dux* TKO ESCs (Supplementary Figure 2C). These luciferase assay results were consistent with our RNA-seq data (Supplementary Figure 2E).

We next performed RNA-seq to identify genes induced by *Ctbp1/2* loss that are DUX-dependent or DUX-independent. A scatter plot comparing gene expression changes between WT and *Ctbp1/2* DKO (Y-axis) and between *Ctbp1/2/Dux* TKO and *Dux* KO (X-axis) revealed a moderate correlation (R = 0.52), suggesting that a substantial population of CtBP1/2-regulated genes act independent of DUX (Figure 2D). Venn diagram showed that more than 75% of genes upregulated in DKO were also upregulated in TKO ESCs (Figure 2E).

To classify these regulatory patterns, we applied K-means clustering analysis (K=5) using the 2,000 most variable genes across WT, *Dux* KO, *Ctbp1/2* DKO, and *Ctbp1/2/Dux* TKO ESCs (Figure 2F, G). Cluster 1 genes (e.g., *Zscan4*, *Usp17*) were slightly repressed in *Dux* KO ESCs but strongly upregulated in *Ctbp1/2* DKO ESCs, and this upregulation was abolished by *Dux* KO, defining DUX-dependent minor ZGA genes. Cluster 2 genes (e.g., *Gadd45g*) were upregulated in both *Ctbp1/2* DKO and *Ctbp1/2/Dux* TKO ESCs, representing DUX-independent target. Cluster 3 genes (e.g., *Gata1*, *Pramel7*, *Zscan5b*) showed moderate downregulation in *Dux* KO ESCs and were modestly upregulated in both *Ctbp1/2* DKO and *Ctbp1/2/Dux* TKO ESCs, corresponding to major ZGA and maternal genes. Cluster 4 (e.g, *Spic*) genes were downregulated by *Dux* KO in WT ESCs. However, this downregulation by *Dux* KO was not observed in *Ctbp1/2* DKO ESCs, indicating that the loss of CtBP1/2 abolishes the dependency of these genes on DUX. Cluster 5 genes were consistently downregulated in both *Ctbp1/2* DKO and *Ctbp1/2/Dux* TKO ESCs, indicating positive regulation by CtBP1/2 in a DUX-independent manner. To evaluate the expression dynamics of these clusters in early mouse embryos, we compared these clusters with public RNA-seq data from early embyos (Zhang et al., 2016), excluding genes with an FPKM average ≤ 1 (Figure 2H). Cluster 1 genes were transient activated at the 2-4-cell stage, typical of minor ZGA genes, whereas Cluster2-4 included maternal genes downregulated post-fertilization and zygotic genes activated after the late 2-cell stage.

Finally, to further examine the dependency of genes upregulated by *Ctbp1/2* DKO on DUX, we measured the expression changes of genes belonging to Cluster 1 and 3 upon inducible *Dux* expression using doxycycline-inducible *Dux* ESCs (Figure 2I). Consistent with our RNA-seq data, Cluster 1 genes were consistently upregulated whereas Cluster 3 genes showed little or no change following short-term DUX induction. Taken together, these results demonstrate that the *Ctbp1/2* depletion broadly upregulates DUX-dependent minor ZGA genes and DUX-independent maternal and major ZGA genes, revealing a dual regulatory system that links CtBP1/2-mediated repression to early embryonic transcriptional programs.

### Temporal regulation of ZGA-associated genes by CtBP1/2 defines rapid and delayed response genes

To monitor the dynamics of ZGA-asscoated gene expression in response to CtBP2 induction and depletion in ESCs, we established *Ctbp1/2* DKO ESCs carrying a doxycycline-inducible CtBP2 transgene. Using this system, we analyzed the expression of representative ZGA-related genes by qRT-PCR. *Dux*, MERVL, *Zscan4c* and *Zfp352* were markedly downregulated between day 3 and day 9 following CtBP2 induction, and subsequently reactivated from day 12 to day 18 upon CtBP2 depletion (Figure 3A). In contrast, *Pramel7* exhibited a rapid response, being strongly downregulated between day 0 and day 3, and then upregulated between day 9 and day 12.

**Figure 3.**
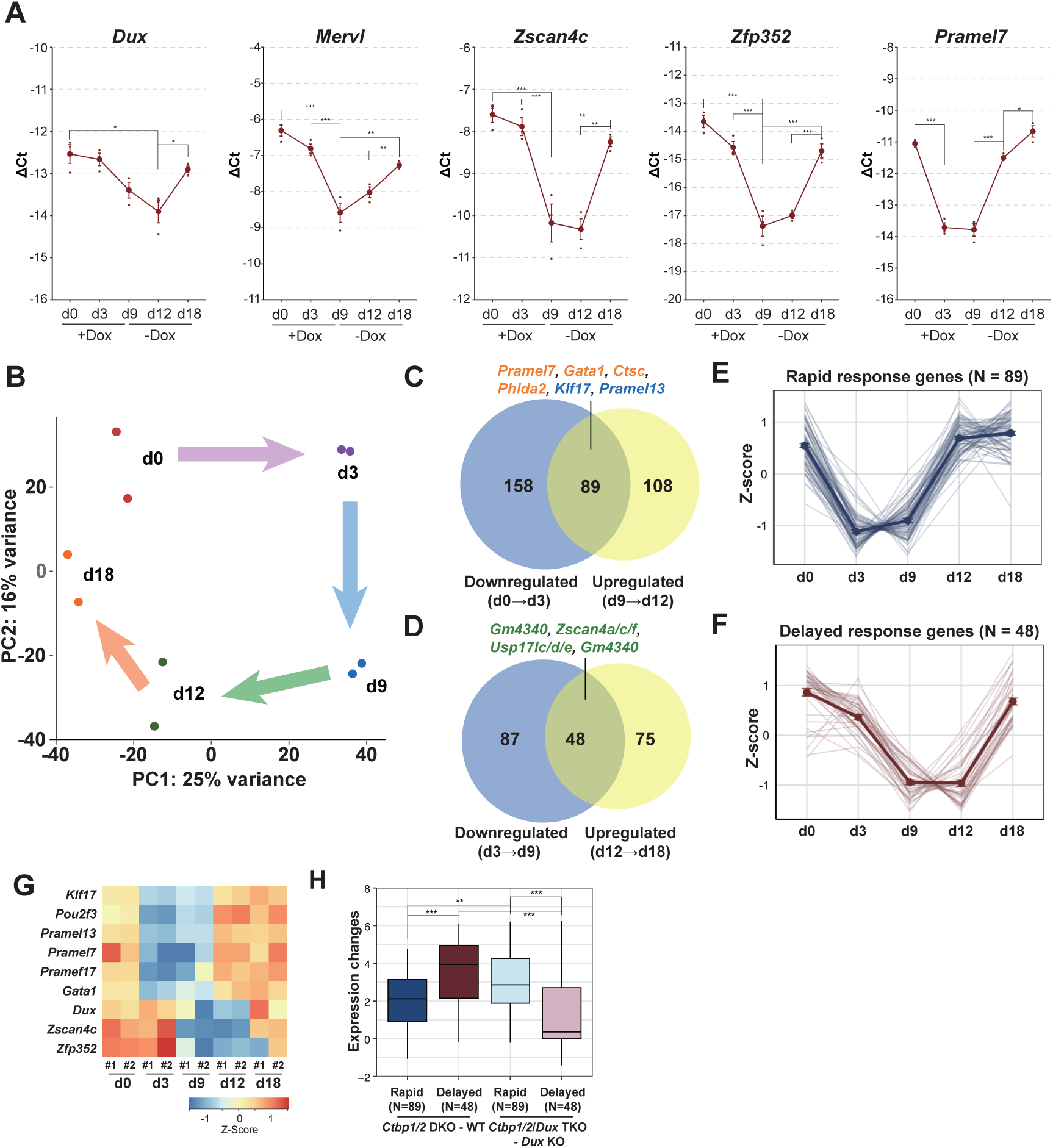
Temporal regulation of ZGA genes by CtBP1/2 defines rapid and delayed responses. (A) RT-qPCR analysis of *Dux*, MERVL*, Zscan4c Zfp352* and *Pramel7* in *Ctbp1/2* DKO ESCs with or without doxycycline (Dox). The ΔCt of each gene was calculated by subtracting the average Ct of *Gapdh* from that of each gene Ct. Means of ΔCt were calculated from three biological triplicates. Error bars indicate the standard error of means (SEM). \*\*\**P* < 0.001, \*\**P* < 0.005, \**P* < 0.05 (Tukey-Kramer multiple comparison test). (B) Principal component analysis (PCA) of *Ctbp1/2* DKO ESCs with or without Dox. (C) Venn diagram showing the overlap between genes downregulated 3 days after CtBP2 induction and genes upregulated 3 days after CtBP2 depletion. (D) Venn diagram showing the overlap between genes downregulated during CtBP2 induction (day3-9) and genes upregulated during CtBP2 depletion (day 12-18). (E, F) Temporal expression dynamics of rapid and delayed response genes upon CtBP2 induction or depletion. Individual gene trajectories (thin lines) and the mean ± SEM (bold lines with error bars) are shown for Rapid response genes (N = 89, panel E) and Delayed response genes (N = 48, panel F). (G) Heatmap showing the expression profiles of selected rapid and delayed response genes. (H) Boxplot showing expression changes of rapid and delayed response genes upon CtBP1/2 depletion in wild-type and *Dux* KO backgrounds.

To distinguish between rapid and delayed transcriptional responses, we performed RNA-seq using *Ctbp1/2* conditional KO (cKO) ESCs at defined time points. Principal component analysis (PCA) revealed that short-term CtBP2 induction (days 0-3) and depletion (days 9-12) drove opposing changes along the PC1 axis, whereas long-term induction (days 3-9) and depletion (days 12-18) contributed to opposite shifts along PC2 (Figure 3B). By overlapping differentially expressed genes under these conditions, we identified two kinetically distinct groups: rapid-response genes, including major ZGA genes (*Pramel7*, *Gata1*, *Phlda2*) and maternal factors (*Klf17*, *Pramel13*) (Figure 3C), and delayed-response genes, comprising minor ZGA genes such as *Zscan4*, *Usp17l*, and *Gm4340* (Figure 3D). Temporal trajectory analysis showed that rapid-response genes (N = 89) were immediately repressed upon CtBP2 induction (days 0-3) and quickly reactivated following CtBP2 depletion (days 9-12) (Figure 3E). By contrast, delayed-response genes (N = 48) displayed gradual downregulation during CtBP2 expression and delayed recovery upon CtBP2 depletion (Figure 3F). Heatmap analysis illustrated their distinct temporal expression patterns (Figure 3G).

Finally, to assess the DUX dependency of these gene sets, we compared gene expression changes induced by *Ctbp1/2* DKO in WT and *Dux* KO backgrounds. Rapid-response genes were equivalently upregulated in both context, whereas the induction of delayed-response genes were significantly attenuated in the absence of DUX (Figure 3H). Collectively, these results indicate that CtBP1/2 represses two transcriptional cascades with distinct temporal kinetics: a DUX-independent program directly targeting major ZGA and maternal genes, and a DUX-dependent program that controls minor ZGA genes.

### CtBP2 directly represses major ZGA genes Pramel7 and Gata1 in ESCs

To examine whether CtBP2 directly regulates major ZGA genes in ESCs, we analyzed the CTBP2 occupancy at rapid- and delayed-response genes using public ChIP-seq datasets (Kim et al., 2015). Genomic peaks were annotated to nearby genes using Genomic Regions Enrichment of Annotations tools (GREAT) with the “two nearest genes” setting (McLean et al., 2010), which assigns each regulatory region to the two closest transcription start sites (TSSs) within 1 Mb (Figure 4A). Approximately half of the rapid-response genes (40 out of 89), but only 2 of 48 delayed-response genes, contained CtBP2-binding peaks. Genome browser views of H3K27Ac (Zhang et al., 2020), a marker for active enhancers, DUX (Hendrickson et al., 2017), and CtBP2 occupancy demonstrated that CtBP2 was bound at regulatory regions of rapid-response genes such as *Pramel7* and *Gata1*, whereas DUX binds to delayed-response loci such as *Duxf3* and *Zscan4c* (Figure 4B, C).

**Figure 4.**
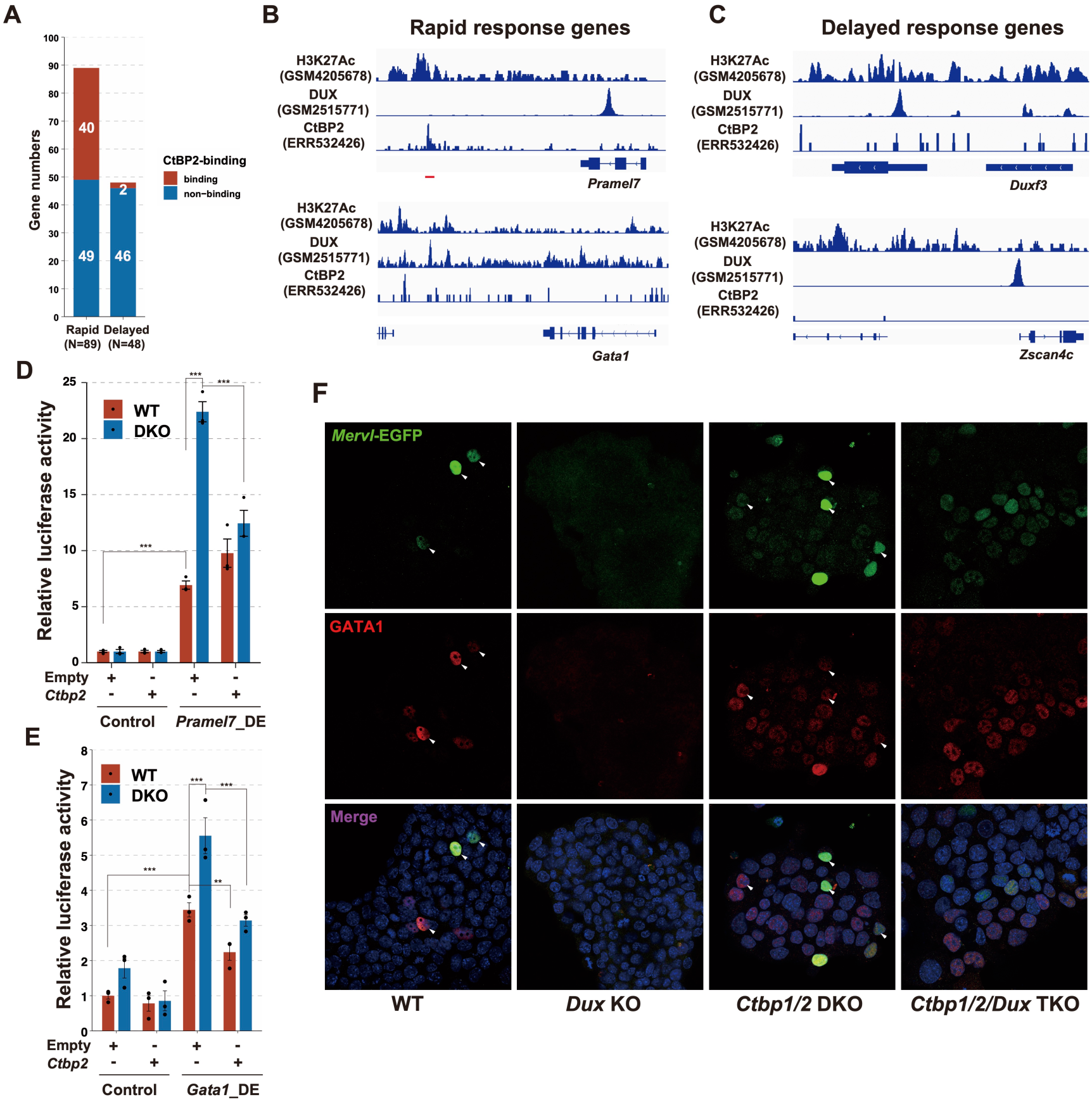
Major ZGA genes, *Pramel7* and *Gata1,* are direct targets of CtBP2 in ESCs. (A) Proportion of CtBP2-bound genes in rapid- and delayed-response groups based on public CtBP2 ChIP-seq data (Kim et al., 2015). Peak-to-gene assignment performed using GREAT (McLean et al., 2010) under the “two nearest genes” rule (≤1 Mb). (B, C) Genome browser views showing H3K27ac (Zhang et al., 2020), DUX (Hendrickson et al., 2017) and CtBP2 ChIP-seq signals at representative rapid (B) and delayed (C) response genes. (D, E) Luciferase assays using distal regulatory regions of *Pramel7* (D) or *Gata1* (E) loci in wild-type (WT) and *Ctbp1/2* DKO ESCs with or without CtBP2 overexpression. (F) Immunofluorescence analysis of WT, *Dux* KO, *Ctbp1/2* DKO and *Ctbp1/2/Dux* TKO ESCs stained with anti-GFP (green) and GATA1 (red) antibodies. Nuclei were counterstained with DAPI. Arrowheads indicate MERVL-EGFP-positive cells.

To functionally validate the repressive activity of CtBP2-bound regions, we performed luciferase reporter assays using distal enhancer fragments derived from *Pramel7* and *Gata1* loci in wild-type and *Ctbp1/2* DKO ESCs. Each enhancer activities was elevated in *Ctbp1/2* DKO ESCs and was suppressed upon CtBP2 re-expression (Figure 4D, E), supporting the idea that CtBP2 directly represses *Gata1* and *Pramel7* transcription through their distal enhancer elements in ESCs.

Immunofluorescence analysis further showed that MERVL-EGFP-positive cells expressed GATA1 in WT ESCs, whereas this expression was abolished in *Dux* KO ESCs (Figure 4F). Consistent with the FACS profiles described earlier (Figure 2A), we observed both EGFP-weak and strong-populations in *Ctbp1/2* DKO ESCs, and both populations expressed GATA1. Deletion of *Dux* in *Ctbp1/2* DKO ESCs abolished selectively eliminated the EGFP-strong, and GATA1-positive cells, whereas the EGFP-weak-positive cells, GATA1-positive cells persisted. These findings suggest that *Ctbp1/2* DKO ESCs comprise two transcriptionally distinct 2CLC population: (1) a DUX-dependent, GATA1-positive subset corresponding to classical 2CLCs marked by strong MERVL-EGFP-strong expression, and (2) a DUX-independent, GATA1-positive subset representing a novel 2CLC state associated with major ZGA gene activation.

### PRAMEL7 regulates subsets of both DUX-dependent and -independent ZGA-associated genes

*Pramel7* belongs to the PRAME (preferentially expressed antigen in melanoma) gene family, whose expression is normally restricted to testis and cancers (Kern et al., 2021). As the third largest gene family in mouse genome (Church et al., 2009), we examined its regulation in wild-type (WT), *Dux* KO, *Ctbp1/2* DKO and *Ctbp1/2/Dux* TKO ESCs. A phylogenetic tree constructed from amino acid sequences annotated in mm39 revealed that PRAME family genes are organized into six chromosomal clustres, with high intracluster similarity (Supplementary Figure 3A, B). RNA-seq analysis showed that most PRAME—except *Lrrc14*, *Lrrc14b*, and *Pramel48*—were markedly upregulated in *Ctbp1/2* DKO ESCs (Figure 5A).

**Figure 5.**
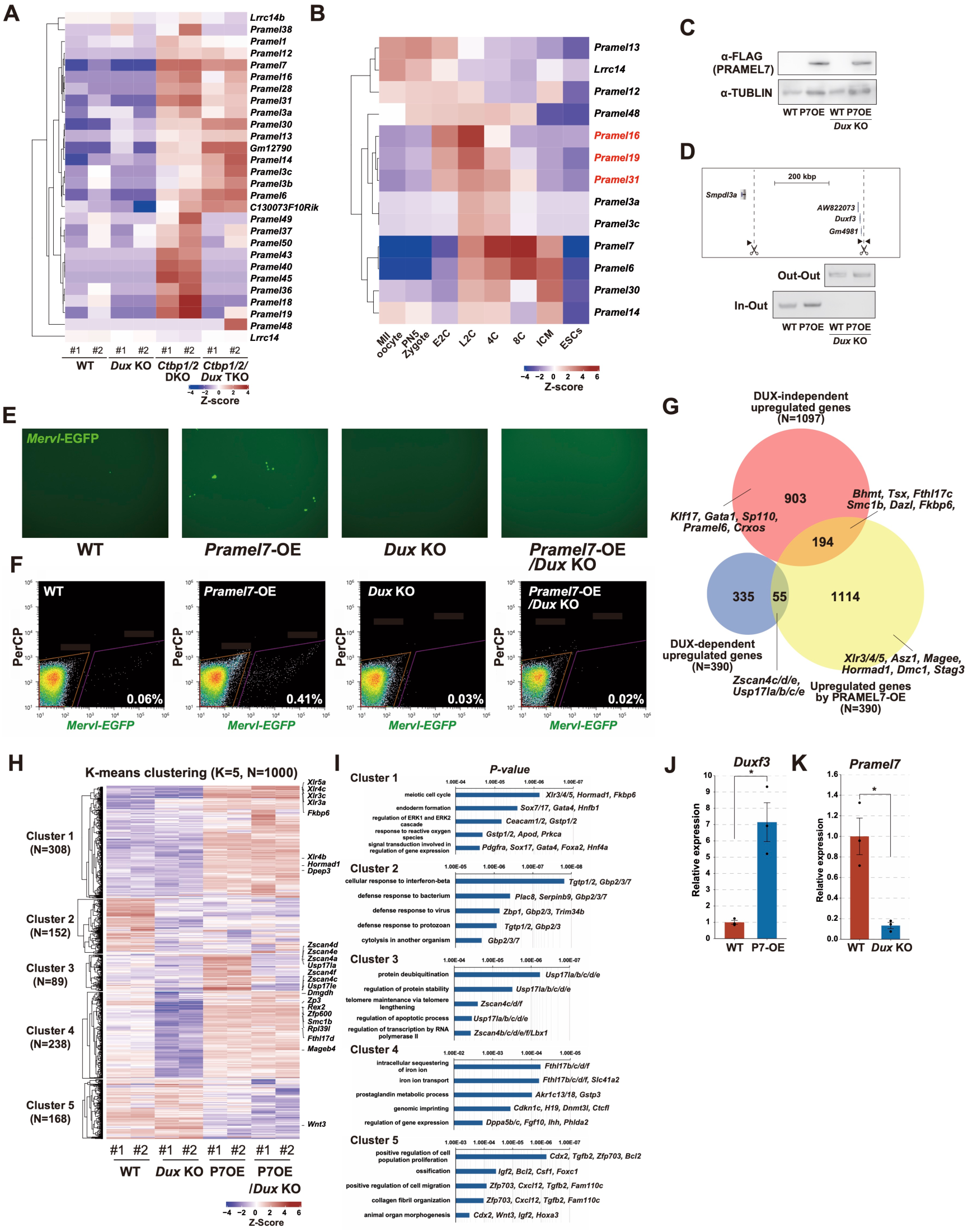
PRAMEL7 controls a subset of ZGA-associated genes through DUX-dependent and -independent mechanisms. (A) Hierarchical clustering heatmap of of PRAME family genes from RNA-seq of wild-type (WT), *Dux* KO, *Ctbp1/2* DKO and *Ctbp1/2*/*Dux* KO ESCs. (B) Expression dynamics of PRAME family genes during early mouse embryos (Zhang et al., 2016). Red letters indicate DUX-dependent genes. (C) Western blot analysis of FLAG-tagged PRAMEL7 protein levels WT, PRAMEL7-overexpressing (P7OE), *Dux* KO and P7OE/*Dux* KOESCs. (D) Genotype PCR confirming large deletion of *Dux* cluster locus using primer sets flanking the gRNA target site. (E) Fluorescence images showing MERVL-EGFP signals in WT, P7OE, *Dux* KO and P7OE/*Dux* KO ESCs. (F) FACS analysis quantifying the percentage of MERVL-EGFP-positive cells in WT, P7OE, *Dux* KO and P7OE/*Dux* KO ESCs. (G) Venn diagram showing the overlap of genes upregulated by *Ctbp1/2* DKO (with or without DUX) and those induced by PRAMEL7 overexpression (H) K-means clustering (K=5) of the top 1,000 most variable genes from RNA-seq of WT, P7OE, *Dux* KO and P7OE/*Dux* KO ESCs. ((I) Gen ontology enrichment analysis of each gene cluster identified in (H). (J) RT-qPCR analysis of *Duxf3* in WT and P7OE ESCs. (K) RT-qPCR analysis of *Pramel7* in WT and *<u>Dux</u>* KO ESCs. The ΔCt values were calculated relative to *Gapdh*, and error bar indicates the standard error of means (SEM) from three biological replicates. Statistical significance was assessed by Tukey-Kramer multiple comparison test (\*\*\**P* < 0.001, \*\**P* < 0.005, \**P* < 0.05).

Based on expression patterns across WT, *Dux* KO, *Ctbp1/2* DKO and *Ctbp1/2/Dux* TKO ESCs, PRAME genes were classified into four groups: (1) genes unaffected by *Ctbp1/2* DKO (e.g., Lrrc14); (2) genes further upregulated by *Dux* KO (e.g., *Pramel14/30*); (3) genes suppressed by *Dux* KO (e.g., *Pramel16/19/31*, DUX-dependent); and (4) genes unaffected by *Dux* KO (e.g., *Pramel7/12*, DUX-independent). Single-cell RNA-seq analysis of early embryonic stages (Zhang et al., 2016) revealed that DUX-dependent PRAME genes (e.g., *Pramel16/19/31*) are transiently induced at the mid-2-cell stage. In contrast, DUX-independent PRAME genes could be further subdivided into two categories (Figure 5B): (i) *Pramel6/7/30*, which are upregulated from mid-2-cell and peak at ICM before being silenced in ESCs; and (ii) *Pramel12/13*, which are maternally expressed and downregulated by at 4-cell stage.

Given the DUX-independent function of *Pramel7* and its known importance in early mouse development (Graf et al., 2017), we generated ESCs overexpressing *Pramel7* (P7OE) in both WT and *Dux* KO backgrounds (Figure 5C, D). *Pramel7* overexpression increased the fraction of MERVL-EGFP-positive 2CLCs (Figure 5E, F), although to a lesser extent than in *Ctbp1/2* DKO. This increase was abolished in *Dux* KO background, and the MERVL-EGFP-weak positive population was absent, indicating that PRAMEL7-driven 2CLC partially depends on DUX.

We next compared genes upregulated by PRAMEL7 with those upregulated by *Ctbp/1/2* DKO in either DUX-dependent or -independent contexts. Venn diagram analysis revealed that a subset of genes activated in a DUX-dependent (55 out of 390) as well as DUX-independent manner (194 out of 1,097) upon *Ctbp1/2* DKO were also upregulated by PRAMEL7 overexpression (Figure 5G). To further investigate the transcriptional programs controlled by PRAMEL7, we performed K-means clustering (K=5) of the top 1,000 most variable genes from RNA-seq of WT, *Dux* KO, P7OE and P7OE/*Dux* KO ESCs (Figure 5H). Cluster 1 contained PRAMEL7-induced genes unaffected by *Dux* KO, including germline-specific and DNA methylation-repressed genes (e.g., *Xlr3/4/5*, *Hormad1*) (Escalier et al., 2002; Shin et al., 2010), which were enriched for meiotic cell cycle pathways (Figure 5I). Cluster 3 genes, including *Zscan4* and *Usp17l* family members, were upregulated by PRAMEL7 overexpression but completely abolished upon *Dux* KO background (Figure 5J), indicating that PRAMEL7 indirectly promotes the expression of minor ZGA genes through *Dux* activation, consistent with increased *Duxf3* expression (Figure 5J). Cluster 4 comprised genes downregulated by *Dux* KO ESCs in WT ESCs but not in P7OE ESCs, suggesting that these genes are suppressed as a consequence of *Pramel7* downregulation in the absence of DUX. Consistent with this, *Pramel7* expression was significantly reduced in Dux KO ESCs (Figure 5K). Together, these results identify PRAMEL7 as a key mediator that links DUX-dependent and -independent transcriptional programs in establishing the totipotent-like states.

### Dux controls part of the ZGA transcriptional program through PRAMEL7 functions

To investigate the functional relationship between DUX and PRAMEL7, we generated *Pramel7* KO ESCs and compared transcriptomic changes between *Dux* KO and *Pramel7* KO ESCs using scatter plot analysis (Figure 6A). Global expression alterations showed a moderate positive correlation (*R* = 0.39), particularly among downregulated genes. Gene ontology (GO) analysis of genes downregulated in both *Pramel7* KO and *Dux* KO ESCs revealed enrichment for terms related to “Nucleic acid binding” and “Homeodomain”, including *Zfp600*, *Zic2* and *Cphx*, *Pou3f1*. In contrast, genes downregulated genes only in *Dux* KO were enriched for “Multicellular organism development” and “Metal ion binding”, including *Piwil1*, *Dppa5* and *Zsacn4c*, *Gata1/2* (Figure 6B). The Overlap between genes upregulated in *Dux* KO and *Pramel7* KO was modest compared with strong overlap observed for downregulated genes (Figure 6C). Furthermore, GSEA demonstrated that the overexpression of PRAMEL7 in *Dux* KO ESCs effectively rescued the downregulated genes resulting from *Dux* KO (Figure 6D), indicating that downregulation of *Pramel7* expression is major cause of the transcriptional abnormalities in *Dux* KO ESCs.

**Figure 6.**
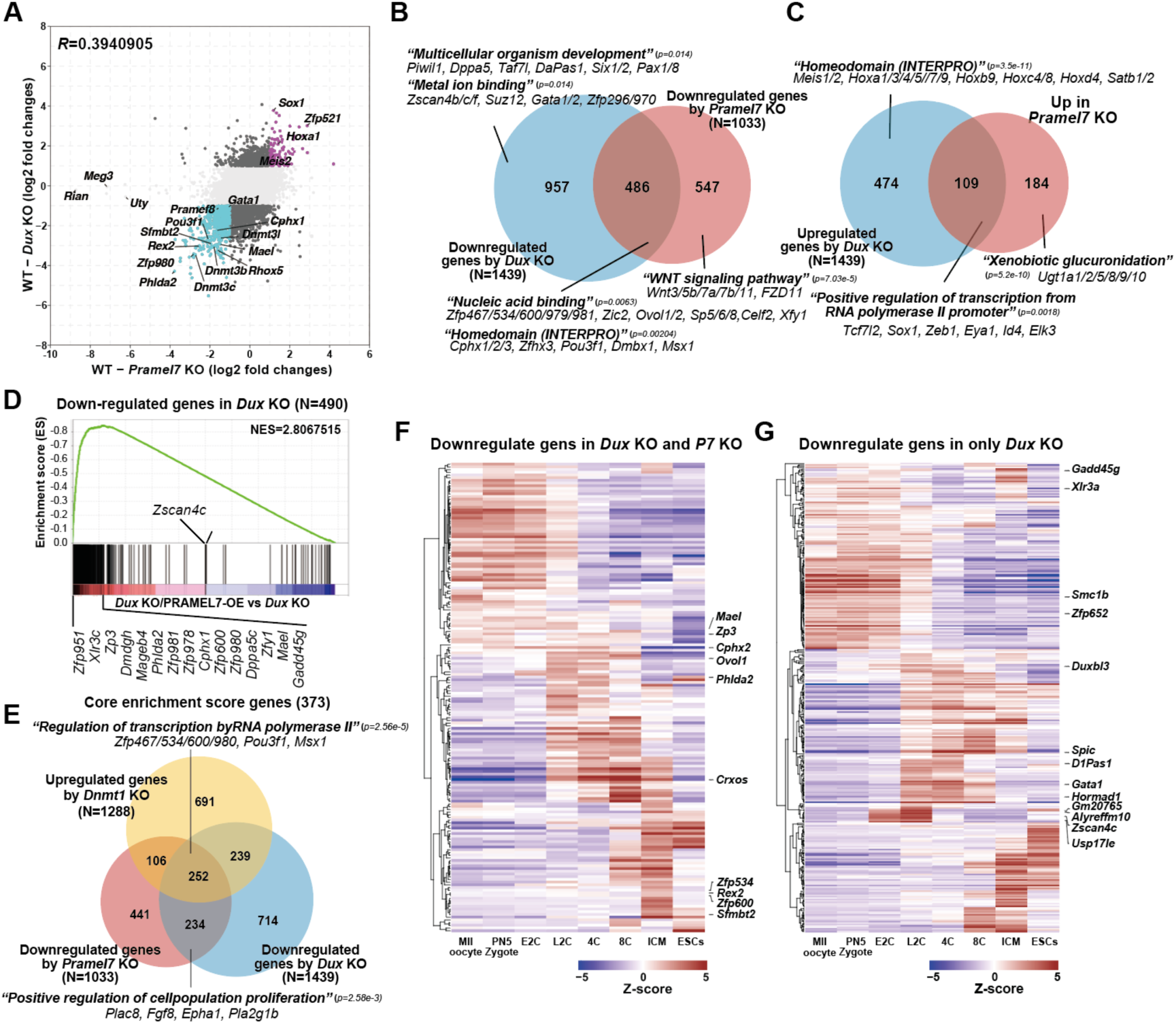
DUX regulates part of ZGA gene program through activation or *Pramel7* in ESCs. (A) Scatter plot showing the correlation between expression changes in *Dux* KO and *Pramel7* KO ESCs. (B) Venn diagram showing the overlapping of the downregulated genes by *Dux* KO in ESCs with the downregulated genes by *Pramel7* KO in ESCs. (C) Venn diagram showing the overlapping of the upregulated genes by *Dux* KO in ESCs with the upregulated genes by *Pramel7* KO in ESCs. (D) Gene set enrichment analysis (GSEA) using genes downregulated by the *Dux* KO to compare gene expression between PRMAEL7-overexpressing in *Dux* KO ESCs and *Dux* KO ESCs. (E) Venn diagram showing the overlap among genes upregulated in *Dnmt1* KO ESCs and genes downregulated in *Pramel7* KO and *Dux* KO ESCs. (F, G) Heatmaps showing hierarchical clustering of genes downregulated genes in both *Dux* KO and *Pramel7* KO ESCs (F), or exclusively in *Dux* KO ESCs (G), using public RNA-seq data of early mouse embryo (Zhang et al., 2016).

**Figure 7.**
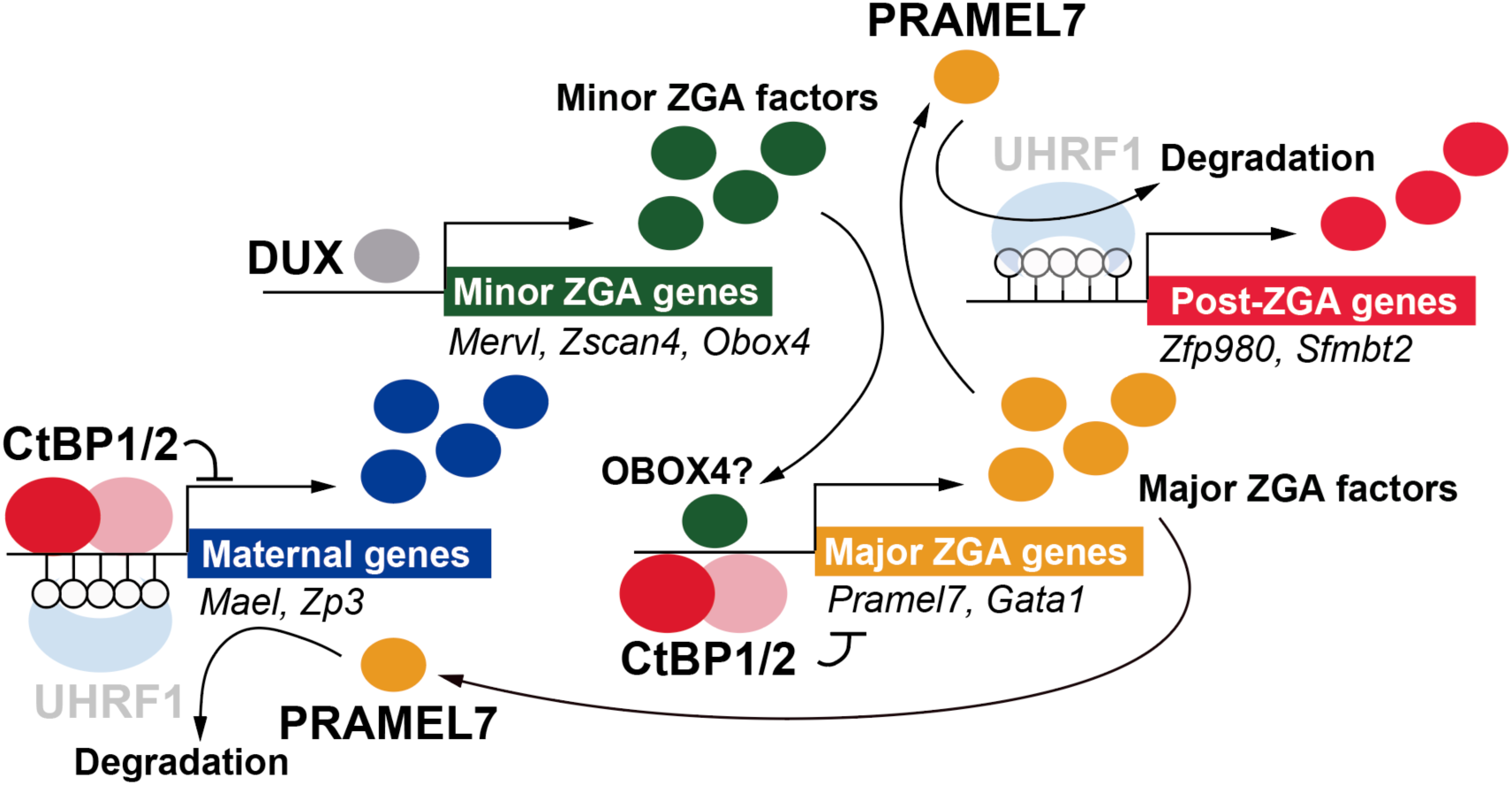
A schematic model for the regulation of 2C-related genes by DUX, CtBP1/2 and PRAMEL7. In this study, we demonstrate that the loss of *Ctbp1/2* in ESCs activates both DUX-dependent and DUX-independent transcriptional cascades to establish a totipotent-like state (Figure 7). Most DUX-dependent genes activated upon *Ctbp1/2* depletion correspond to minor ZGA genes transiently expressed at the two-cell (2C) stage. In contrast, DUX-independent genes upregulated upon *Ctbp1/2* loss can be classified into maternally expressed genes and post-major ZGA genes. Among them, *Pramel7* is directly upregulated by *Ctbp1/2* loss in a DUX-independent manner and further modulates both DUX-dependent and -independent pathways through the activation of DUX and its downstream targets. Collectively, these findings uncover a dual regulatory framework in which CtBP1/2 represses both DUX-mediated and DUX-independent transcriptional programs, providing new insights into molecular mechanisms governing early mouse embryogenesis.

Because PRAMEL7 is known to promote UHRF1 degradation, we next examined whether DNA methylation contributes to gene silencing in *Dux* and *Pramel7* KO ESCs. More than half of the genes downregulated in both *Dux* KO and *Pramel7* ESCs were upregulated by *Dnmt1* KO ESCs (Figure 6E), suggesting that the transcriptional repression observed in these mutants is mediated, at least in part, by aberrant DNA methylation.

We next analyzed the expression dynamics of genes downregulated in both *Dux* KO and *Pramel7* KO ESCs, as well as those specifically downregulated only in *Dux* KO ESCs, using public single-cell RNA-seq data from early mouse embryos (Zhang et al., 2016). Gene downregulated genes both in *Dux* KO and *Pramel7* KO ESCs were primarily associated with maternally expressed genes, major ZGA genes and post-ZGA genes (Figure 6F). In contrast, genes downregulated only in *Dux* KO ESCs included not only maternally expressed genes, major ZGA genes and post-ZGA genes but also minor ZGA genes (Figure 6G).

Taken together, these findings suggest a mechanistic model in which DUX primarily activate minor ZGA genes, whereas PRAMEL7―one of the indirect downstream targets of DUX―regulates maternally, major ZGA genes and post-ZGA genes through modulation of DNA methylation mediated by UHRF1 degradation in 2CLCs.

## Discussion

Our data clearly demonstrate that the loss of *Ctbp1/2* in ESCs upregulates maternally expressed and post-major ZGA genes in a DUX-independent manner, highlighting an alternative regulatory axis operating during early embryonic development. Interestingly, recent studies have demonstrated that OBOX proteins, a family of PRD-like homeobox transcription factors, can compensate for DUX function by directly activating MERVL as well as both minor and major ZGA genes (Guo et al., 2024; Ji et al., 2023; Sakamoto et al., 2024). However, our transcriptome analysis revealed that only *Obox4* was upregulated in *Ctbp1/2* double-knockout (DKO) ESCs, and this upregulation was completely abolished by *Dux* KO. These results suggest that the loss of *Ctbp1/2* regulates early developmental genes through pathways independent of both DUX and OBOX.

By integrating transcriptome data from conditional *Ctbp2* KO ESCs with public CtBP2 ChIP-seq datasets, we identified direct CtBP2 target genes (e.g., *Pramel7*, *Gata1*, and *Klf17*) and distinguished them from DUX-dependent regulatory genes. Furthermore, we found that MERVL-EGFP-weak positive cells in both *Ctbp1/2* DKO and *Ctbp1/2/Dux* TKO ESCs expressed GATA1, a direct CtBP2 target. Importantly, GATA1-positive cells were absent in *Dux* KO ESCs, suggesting that *Ctbp1/2* loss not only relieves DUX dependency but also rewires the transcriptional hierarchy governing ZGA-associated programs.

*Prame* is one of the most amplified gene families in mammals, and the mouse *Prame* gene family represents the third-largest multigene family in the genome (Chang et al., 2011). We showed that the disruption of *Ctbp1/2* activates the majority of *Prame* family genes in ESCs, some of which are upregulated by DUX, while others are induced directly by the loss of CtBP1/2. DUX-dependent *Prame* genes are enriched among minor ZGA genes, whereas DUX-independent *Prame* genes are upregulated after major ZGA. Moreover, many *Prame* genes are rapidly repressed upon the re-expression of CtBP2 in *Ctbp1/2* DKO ESCs, indicating direct repression by CtBP1/2.

The positive regulators of *Prame* family genes remain largely unknown. Notably, published RNA-seq data indicate that OBOX3/5, but not DUX, can activate *Pramel7* and *Pramef17* (*Pramel14*), both of which are upregulated by *Ctbp1/2* loss even in the absence of DUX. In contrast, *Pramel16*, *Gm12794* (*Pramel19*) and *A430089l19Rik* (*Pramel39*) require DUX for activation (Sakamoto et al., 2024). Because *Obox* family genes were silenced in *Ctbp1/2/Dux* TKO ESCs, it is possible that another positive regulator, distinct from OBOX proteins, is activated upon *Ctbp1/2* depletion or constitutively expressed in WT ESCs. Expression of *Prame* family genes has been observed in various cancer and germline cells (Kern et al., 2021), and their expression pattern resembles that of *Pou5f1* (Oct4) (Bortvin et al., 2003), suggesting that an OCT4-centered transcriptional network may function as a positive regulatory axis for *Prame* genes.

We further demonstrated that the overexpression of PRAMEL7 partially rescues the transcriptional abnormalities observed in *Dux* KO ESCs, in which *Pramel7* expression was markedly reduced. PRAMEL7 activates two distinct genetic programs: one involving minor ZGA genes through *Dux* upregulation, and another encompassing maternally expressed, major ZGA and post-ZGA genes. Previous work showed that PRAMEL7 promotes global DNA demethylation by targeting UHRF1 for proteasomal degradation in ESCs (Graf et al., 2017). In addition, *Dux* expression is epigenetically repressed by DNA methylation maintained by the ZBTB24-CDCA7-HELLS axis (Guo et al., 2025). These findings suggest that PRAMEL7 regulates DUX-dependent transcriptional cascades by promoting DNA demethylation at the *Dux* locus. Furthermore, we found that maternally expressed and major ZGA genes regulated by PRAMEL7 are also activated upon *Dnmt1* depletion, indicating that PRAMEL7 controls these genes through DNA demethylation mediated by UHRF1 degradation. Because PRAMEL7 also promotes degradation of NuRD complex at chromatin to sustain pluripotency in ESCs (Rupasinghe et al., 2024), it will be important to explore whether NuRD degradation by PRAMEL7 contributes to the regulation of ZGA-associated transcriptional programs.

Although DUX is a master regulator of the transition from pluripotent ESCs to 2-cell-like cells, our results demonstrated that PRAMEL7 overexpression rescues most of the defects observed in *Dux* KO ESCs. *Prame* family genes are upregulated from the early phase of 2CLC emergence traced by the *Zscan4* and MERVL reporter system based on scRNA-seq (Eckersley-Maslin et al., 2016; Rodriguez-Terrones et al., 2018), and MERVL-positive 2CLCs are eliminated by *Dux* KO (De Iaco et al., 2017). Our data further showed that *Pramel7* expression was significantly reduced upon *Dux* KO in WT ESCs but remained unaffected in *Ctbp1/2* DKO ESCs, indicating that the loss of *Ctbp1/2* rescues the *Dux* KO phenotypic consequences through *Pramel7* upregulation in ESCs. Taken together, we propose that DUX orchestrates two distinct transcriptional programs in early mouse embryos: (1) direct activation of minor ZGA genes (e.g., MERVL, *Zscan4c* and *Obox4*), and (2) PRAMEL7-mediated transcriptional cascades. While *Dux* KO ESCs exhibit a marked reduction in *Pramel7* expression, *Pramel7* remains normally expressed in *Dux* KO embryos at the mid-2C stage (Guo et al., 2019). OBOX5, which is maternally expressed, represents a strong candidate regulator for the DUX-independent activation of *Pramel7* in *Dux* KO embryos.

Our findings highlight fundamental differences between *in vitro* induction of 2CLCs and *in vivo* regulation of ZGA, which are likely governed by maternal factors such as OBOX and CtBP2. To better characterize these DUX-independent cascades, dual-reporter ESC models (e.g., MERVL-tdTomato/*Gata1* or *Pramel7*-EGFP reporters) combined with single-cell RNA-seq represent powerful approaches to dissect the molecular networks underlying 2CLC heterogeneity induced by *Ctbp/1/2* loss.

## Author Contributions

Y.S. conceived the study and experimental design, performed experiments and wrote the manuscript. K.S., K.Y. and N.H. performed most experiments. R.M. M.M., A.S. and Y.M performed some experiments. K.S., K.Y. and N.H. contributed equally to this work.

## Acknowledgements

We thank all members of Seki laboratory for stimulating discussions. This study was supported by JSPS KAKENHI Grant Numbers JP18H02422, JP20H05375, JP22H04682 and a Special Research Grant for Individual Researchers from Kwansei Gakuin University.

## Data availability

RNA-Seq data has been deposited in Gene Expression Omnibus (GSE253944)

