## Supplementary Figures for "CtBP restricts DUX-dependent and -independent genetic program for the 2-cell-like state in murine embryonic stem cells"

### Supplementary Figure 1

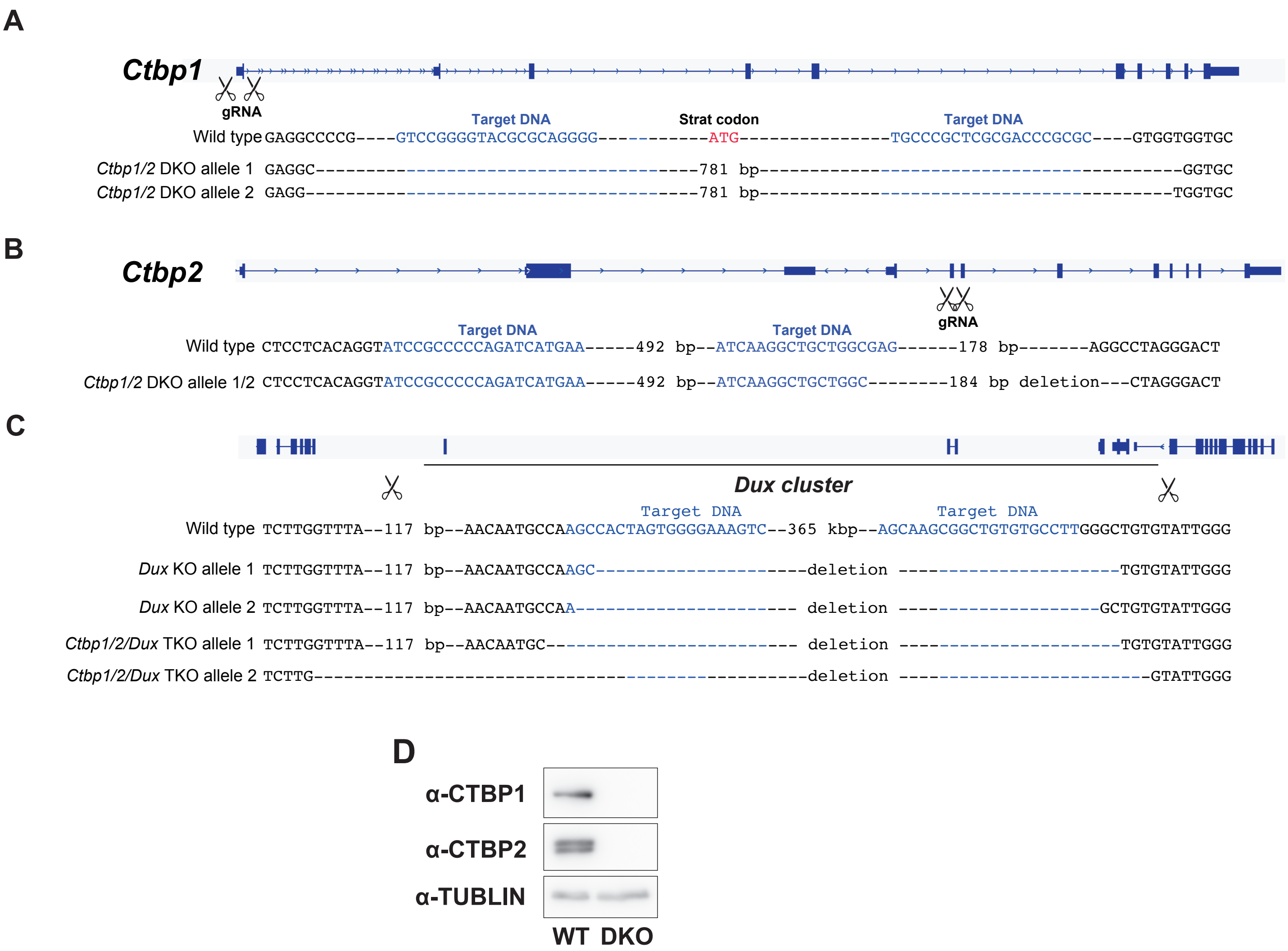


**Supplementary Figure 1.** (A-C) The deletion sequence of *Ctbp1/2* and *Dux* in each KO ESCs. (D) Wesern blotting of CtBP1/2 in WT and *Ctbp1/2* DKO ESCs.

### Supplemantry Figure 2

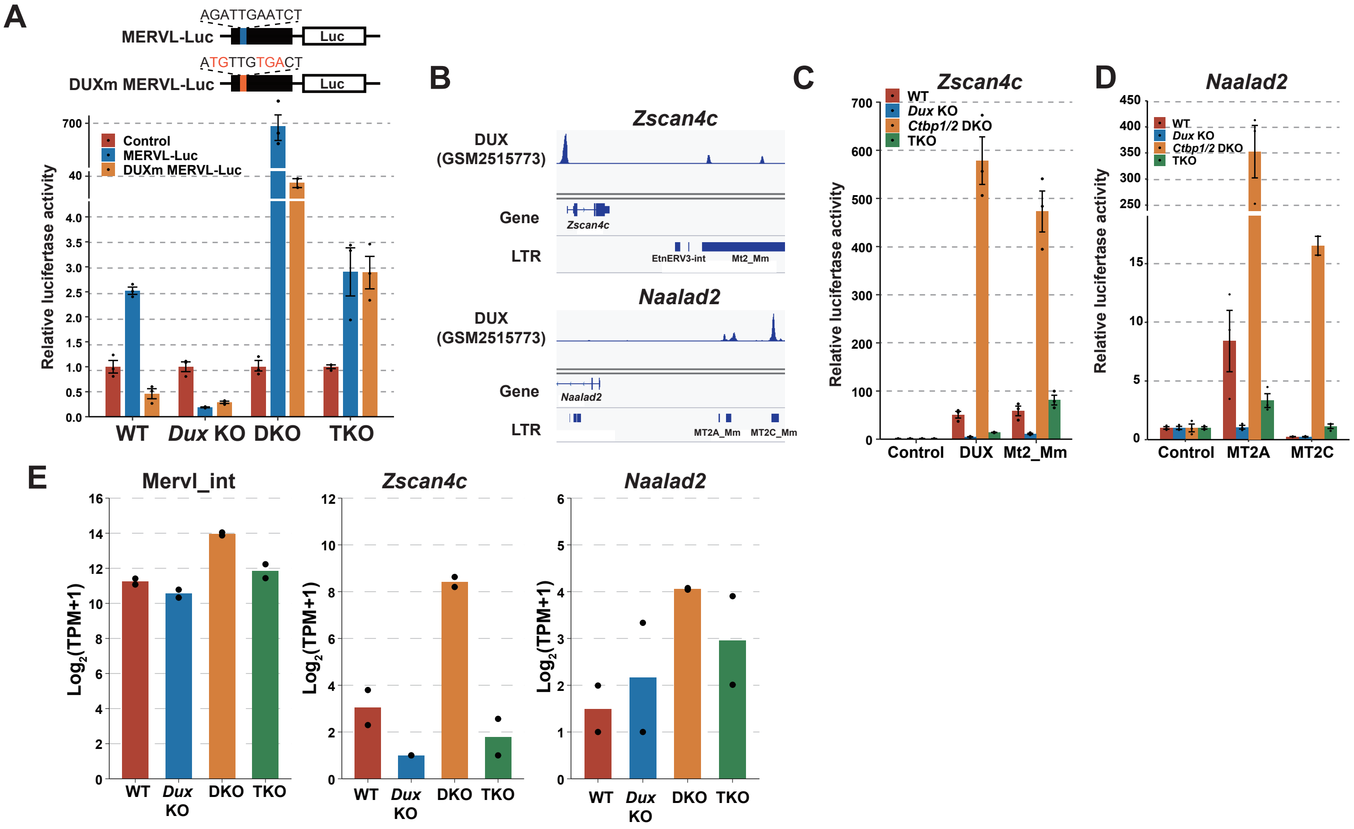


**Supplemantry Figure 2.** (A) Luciferase activity driven by MERVL-LTR or MERVL-LTR with mutations in the DUX recognition sequence in WT, *Dux* KO, *Ctbp1/2* DKO and *Ctbp1/2/Dux* TKO ESCs. (B) Publich ChIP-Seq data of DUX around *Zscan4c* and *Naalad2*. (C, D) Luciferase activity driven by the DUX-binding regions without MERVL or Mt2_Mm around *Zscan4c* (C) or MT2A and MT2C around *Naalad2* (D). (E) Expression levels of Mervl_int, *Zscan4c* and *Naalad2* in WT, *Dux* KO, *Ctbp1/2* DKO and *Ctbp1/2/Dux* TKO. The $\log_2$(TPM+1) values dertermined by two independent RNA-seq analyse are shown.

### Supplementary Figure 3

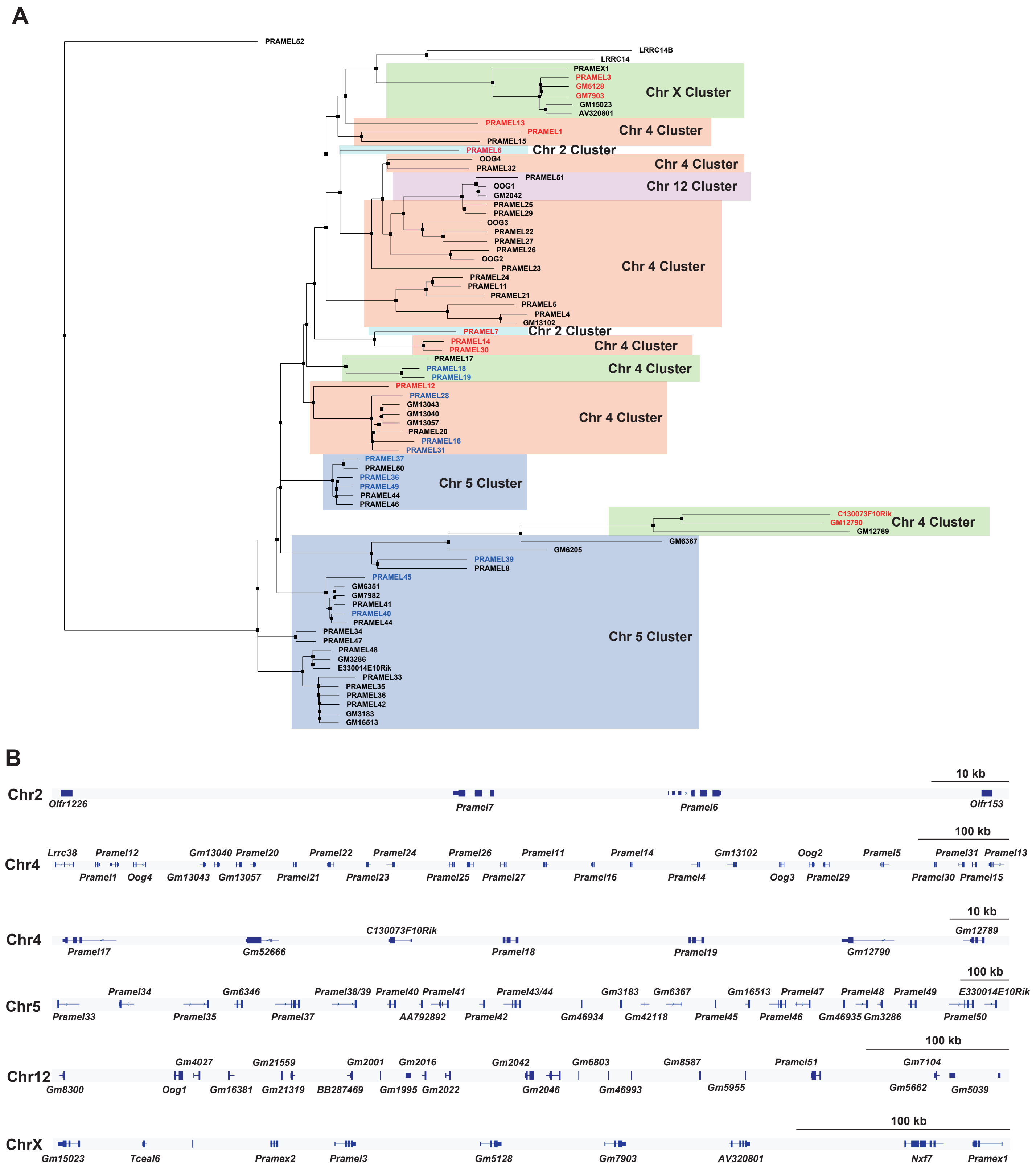


**Supplementary Figure 3. Molecular phylogeny and synteny for PRAME family proteins.** (A) A maximum-likelihood the tree was constructed using amino acid sequences of PRAME family proteins and LRRC14 as an outgroup. Red letters indicate the genes upregulated by *Ctbp1/2* DKO without DUX. Blue letters indicate the genes upregulated by *Ctbp1/2* DKO with DUX. (B) Synteny of Prame family genes at Chromosome 2, 4, 5, 12 and X.
